# Individual variation in the attribution of incentive salience to social cues

**DOI:** 10.1101/582254

**Authors:** Christopher J. Fitzpatrick, Jonathan D. Morrow

## Abstract

Research on the attribution of incentive salience to drug cues has furthered our understanding of drug self-administration in animals as well as drug relapse and craving in humans. The influence of peers and other social cues on drug-seeking has garnered more attention recently, but few studies have investigated the ability of social cues to gain incentive-motivational value. In the present study, a Pavlovian conditioned approach procedure was used to identify rats that are more (sign-trackers) or less (goal-trackers) prone to attribute incentive salience to food reward cues. A novel procedure then employed social ‘peers’ to compare the tendency of sign-trackers and goal-trackers to attribute incentive salience to social reward cues. Social behavior of sign-trackers and goal-trackers was also compared using social interaction and choice tests. Finally, basal levels of plasma oxytocin were measured in sign-trackers and goal-trackers, because oxytocin is known to modulate the mesolimbic reward system and social behavior. Compared to goal-trackers, sign-trackers attributed more incentive salience to social cues and exhibited more prosocial behaviors. No group differences were observed in baseline plasma oxytocin levels. Taken together, these experiments demonstrate a concordance of individual variation in social behavior, the attribution of incentive salience to social cues following peer interaction, and attribution of incentive salience to food cues. This general tendency to attribute motivational value to reward cues has important implications for the pathophysiology of addiction and other disorders of reward learning.

## Introduction

Nearly every neuropsychiatric disorder involves alterations in social behaviors, and several disorders are characterized by abnormal processing of social cues (Adolphs, 2010). Specifically, addiction pathophysiology involves the cooption and alteration of social behaviors and cue processing (Volkow et al., 2011). For example, social interactions within drug-taking contexts enhance drug-seeking behavior (Neisewander et al., 2012), and social cues (e.g., people) can elicit similar reactivity and craving as drug-related cues (Conklin et al., 2013, Larsen et al., 2010). Indeed, in a sample of adolescents, being around other individuals was the greatest contributor to drug relapse (73%) regardless of whether the individuals were using (Roberts et al., 2015). Tobacco and alcohol companies have long exploited social cues to promote drug consumption, incorporating such cues into 42% and 74% of their advertisements, respectively (Schooler et al., 1996). Interestingly, ‘social reinstatement’ of drug-seeking behavior has been recently demonstrated in rats (i.e., social ‘peers’ can serve as discriminative stimuli and increased drug-seeking behavior) (Weiss et al., 2018). However, there remains a lack of research linking the processing of social context and cues with addiction-like behaviors, or determining which individuals may be more or less susceptible to the reward-modifying aspects of social cues (Heilig et al., 2016).

Pavlovian conditioned approach (PCA) procedures have previously been used to investigate individual variation in the attribution of incentive-motivational value to food- and drug-related cues (Robinson and Flagel, 2009, Saunders and Robinson, 2010, Yager et al., 2015, Yager and Robinson, 2015). When environmental cues are paired with rewarding stimuli, some animals (goal-trackers; GTs) use the cue as a predictor of impending reward while others (sign-trackers; STs) attribute incentive-motivational value to the cues, making them reinforcing and capable of motivating behavior even in the absence of the reward itself (Robinson and Flagel, 2009). In addition to increased cue-directed behavior, STs display several other behavioral traits that are believed to contribute to addiction-like behaviors, such as impulsivity (Lovic et al., 2011) and novelty-seeking behavior (Beckmann et al., 2011).

Sign-tracking requires dopamine (DA) activity in the mesolimbic dopamine system and is DA-dependent in the nucleus accumbens (NAc) core. Because both nonsocial and social reward cues are processed through the mesocorticolimbic dopamine system (Bhanji and Delgado, 2014), it is likely that STs would (1) sign-track to social cues and (2) show prosocial behaviors. Although social experience has been shown to modulate sign-tracking behavior (Beckmann and Bardo, 2012, Lomanowska et al., 2011), sign-tracking to nonsexual social cues has not previously been investigated.

To address this, the present study investigated individual variation in (1) the attribution of incentive salience to social cues and (2) social behaviors in rats. In Experiment 1, a novel procedure combining conditioned place and cue preference was used to measure sign-tracking to a social cue as well as social context in GTs and STs. In Experiment 2, social interaction and social choice tests were used to measure to measure sociability and social novelty seeking in GTs and STs. Finally, in Experiment 3, plasma oxytocin (OXT) samples were measured at baseline and levels were correlated with PCA behavior. OXT was measured because it modulates DA release in response to social stimuli (Baskerville and Douglas, 2010) and regulates the salience of social cues (Shamay-Tsoory and Abu-Akel, 2016).

## Experimental procedures

### Animals

Adult male Sprague Dawley rats (250-300 g) were purchased from Charles River Laboratories. Rats were pair-housed and maintained on a 12 h light/dark cycle. Standard rodent chow and water were available *ad libitum*. All procedures were approved by the University Committee on the Use and Care of Animals (University of Michigan; Ann Arbor, MI).

### Apparatus

Modular conditioning chambers (24.1 cm width × 20.5 cm depth x 29.2 cm height; MED Associates, Inc.; St. Albans, VT) were used for Pavlovian conditioned approach training. Each chamber was contained within a sound-attenuating cubicle equipped with a ventilation fan to provide ambient white noise. For Pavlovian conditioning, chambers were equipped with a pellet magazine, an illuminated retractable lever (counterbalanced on the left or right of the pellet magazine), and a red house light on the wall opposite to the pellet magazine. When inserted into the chamber, the retractable lever was illuminated by an LED light within the lever housing. A pellet dispenser delivered banana-flavored food pellets into the pellet magazine, and an infrared sensor inside the pellet magazine detected head entries. For operant conditioning, the lever was removed from the chamber and replaced with two nose-poke ports on either side of the pellet magazine.

A three-chambered apparatus (60 cm width x 90 cm length x 34 cm height; Formtech Plastics; Oak Park, MI) was used for social behaviors. Each chamber (60 cm width x 30 cm length x 34 cm height) consisted of foamboard floors and walls (matte black polyvinyl chloride) and was connected by foamboard dividers with archways (10 cm width x 12 cm height at apex) to allow access between chambers. For the social choice test, the dividers remained in the apparatus. For the social interaction text, all dividers were removed, creating a single open arena (60 cm width x 90 cm length). For the social conditioned place/cue test, two chambers were used, and the third chamber was blocked.

### Pavlovian conditioned approach: procedure

For two days prior to pretraining, rats were familiarized with banana-flavored food pellets (45 mg; Bioserv; Frenchtown, NJ) in their home cages. Twenty-four hours later, rats were placed into the operant chambers and underwent one pretraining session during which the red house-light remained on, but the lever was retracted. Fifty food pellets were delivered on a variable interval (VI) 30 schedule (i.e., one food pellet was delivered on average every 30 s, but actual delivery varied between 0-60 s). All rats consumed all of the food pellets by the end of the pretraining session. Twenty-four hours later, rats underwent daily PCA training sessions over seven days. Each trial during a test session consisted of extension of the illuminated lever (the conditioned stimulus; CS) into the chamber for 8 s on a VI 90 schedule (i.e., one food pellet was delivered on average every 90 s, but actual delivery varied between 30-150 s). Retraction of the lever was immediately followed by the response-independent delivery of one food pellet (the unconditioned stimulus; US) into the pellet magazine. Each test session consisted of 25 trials of CS-US pairings, resulting in a total session length of approximately 40 min. All rats consumed all the food pellets that were delivered.

### Experiment 1: social conditioned place/cue preference test

Rats underwent seven daily PCA training sessions to screen rats as STs, GTs, and IRs. Only STs (n = 12) and GTs (n = 12) were used for further testing. IRs were used as partner rats during conditioning. Seven days after the last session of PCA training, rats were divided into Paired and Unpaired groups (n = 6; phenotype) and underwent a novel social conditioned place/cue preference procedure. The two chambers of the apparatus were differentiated using visual and olfactory cues. The left chamber had a white background with vertical black stripes, was illuminated internally by blue light (DIODER LED light strips; IKEA, Conshohocken, PA; wrapped in blue acetate film), and was wiped with a 1% almond extract solution (Context A).

The right chamber had a white background with black diamonds, was illuminated by yellow light (DIODER LED light strips; wrapped in yellow acetate film), and was wiped with a 0.5% lemon extract solution (Context B). Blue and yellow light was selected, because it has previously been shown that rats can visually discriminate between these colors (Walton and Bornemeier, 1939). Moreover, the internal illumination prevented shadows, which is important as rats tend to remain immobile in shadowed areas of test arenas (Lapiz-Bluhm et al., 2008). In addition, almond and lemon extract were used, because rats do not show a preference or aversion to these neutral, distinguishable odors (Torras-Garcia et al., 2005).

Training consisted of two daily habituation sessions, eight daily conditioning sessions, and two daily test sessions. All sessions were 10 min in length. During habituation sessions, rats were exposed to the apparatus and allowed to freely explore between the two chambers. The average time spent in each chamber over the two habituation sessions was calculated, and the less preferred chamber was selected as the social-paired chamber. During conditioning sessions, a silver foamboard star was used as a discrete cue (star-CS) and placed in the social-but not empty-paired chamber. Use of the star-CS was loosely adapted from a previous study that demonstrated sign-tracking to an ethanol-paired star-CS in mice (Cunningham and Patel, 2007). During conditioning sessions, rats in the paired group were placed on alternating days into an empty chamber or a chamber containing a rat, and access to the unused chamber was blocked. At the start of conditioning, partner rats were weight-matched to subject rats. In addition, rats from the same home cage were never used as partners, and a novel partner rat was used on each conditioning day to prevent decreased social investigation over repeated pairings (Winslow and Camacho, 1995). In the unpaired group, rats were placed on alternating days into one of two empty chambers. Four hours after the sessions, during which paired rats received social interaction and unpaired rats received nothing, paired rats were placed alone in a novel home cage, and unpaired rats received social interaction in a novel home cage. Twenty-four hours after the last conditioning session, rats underwent context and cue tests. During the context test, the star-CS was removed, and rats could freely explore both chambers (Context A and B). During the cue test, the star-CS was placed in the social-paired chamber, but the context in both chambers was changed to a third, neutral context: white walls, illumination by white light, and cleaning with a 70% ethanol solution (Context C). Like the context test, rats could freely explore both chambers.

### Experiment 2: social choice and social interaction tests

Rats underwent seven days of PCA training to screen rats for STs, GTs, and IRs. Only STs (n = 7) and GTs (n = 5) were used for further testing. IRs were used as partner rats during both tests. Seven days after the last session of PCA training, rats underwent a social choice test. The social choice test was conducted in a three-chambered apparatus and consisted of three 10-min phases: habituation, sociability, and social novelty. Each phase immediately followed the previous phase. For all phases, subject rats were initially placed into the middle chamber on the side farthest from the doorways, and chambers were cleaned with a 70% ethanol solution between phases. During the habituation phase, the three-chambered apparatus contained two wire-mesh baskets (23 cm height, 13 cm top diameter, 22 cm bottom diameter) in the left and right chambers. Behavior was recorded during this phase, and a chamber preference was determined for each individual subject rat. During the sociability phase, the non-preferred chamber contained a partner rat inside a wire-mesh basket, and the preferred side contained an empty wire-mesh basket. Placement of partner rats was counterbalanced between left and right chambers. During the social novelty phase, the previously empty basket contained a novel rat, and the now familiar rat was contained within the other basket.

Seven days after the last phase of the social choice test, rats underwent a social interaction test, and novel, weight-matched partner rats were used. Partner rats were placed first in an open arena (60 cm width x 90 cm length), followed shortly after by placement of subject rats in the opposite corner. Like in Experiment 1, partner rats were weight-matched to subject rats and never came from the same home cage as the subject rat. Behavior was recorded by video camera for 10 min, and after the end of each session the arena was cleaned with a 70% ethanol solution.

### Experiment 3: baseline measurement of plasma OXT levels in GTs, IRs, and STs

Rats underwent seven daily PCA training sessions to screen rats as GTs (n = 7), IRs (n = 10), and STs (n = 11). Seven days after the last PCA training session, rats were removed from their home cages and rapidly decapitated. Trunk blood was collected into chilled, EDTA-coated tubes (10 mL BD Vacutainer® tubes; Becton, Dickinson and Company; Franklin Lakes, NJ), and plasma was separated by centrifugation at 1,600 g for 15 min at 4 °C. Next, plasma was aliquoted and stored at −80 °C within 5 min of collection. An enzyme-linked immunosorbent assay (ELISA; ENZO Life Sciences, Inc.; Farmingdale, NY) was used to quantify plasma OXT levels. Unextracted samples were diluted 1:4 in assay buffer as previously described (Kramer et al., 2004, Szeto et al., 2011). The dilution has been shown to reliably fit unextracted samples on a standard curve. Coefficient of variance was used as a cut-off, and three plasma samples were excluded for having a coefficient of variance higher than 20% (ST = 2, IR = 1).

### Statistical analysis

PCA behavior was scored using an index that combines the number, latency, and probability of lever presses (sign-tracking) and magazine entries (goal-tracking) during CS presentations within a session. Briefly, we averaged together the response bias (i.e., number of lever presses and magazine entries for a session; [lever presses – magazine entries] / [lever presses + magazine entries]), latency score (i.e., average latency to perform a lever press or magazine entry during a session; [magazine entry latency – lever press latency]/8), and probability difference (i.e., proportion of lever presses or magazine entries; lever press probability – magazine entry probability). The index scores behavior from +1.0 (absolute sign-tracking) to -1.0 (absolute goal-tracking), with 0 representing no bias (Meyer et al., 2012). Rats were classified using the following cutoffs: STs (x ≥ 0.5), IRs (−0.5 < x < 0.5), and GTs (x ≤ −0.5). Rats were classified by the average PCA index scores from Sessions 6 and 7.

SPSS (Version 24; IBM, Inc.) was used for all statistical analysis. Across PCA training sessions, lever press and magazine entry number, latency, and probability were analyzed using a linear mixed model with an autoregressive (AR1) covariance structure, selected using Akaike’s information criterion (i.e., the lowest number criterion represents the highest quality statistical model using a given covariance structure). In Experiment 1, for the social conditioned place/cue tests, chamber time was analyzed using two-way analysis of variance (ANOVA) with Phenotype (GT and ST) and Group (Unpaired and Unpaired) as factors. In Experiment 2, during the social choice test, chamber time in each phase was analyzed using a two-way ANOVA with Chamber (Left and Right) and Phenotype (GT and ST) as factors. For the social interaction test, number and duration of social interactions as well as number of fecal boli were analyzed using independent samples t-tests with Phenotype (GT and ST) as a factor. In Experiment 3, OXT levels were analyzed using a one-way ANOVA with Phenotype (GT, IR, and ST) as a factor. Correlations were performed using Pearson’s r. With a significant ANOVA, multiple comparisons were performed using Fisher’s Least Significant Difference (LSD) post hoc test.

In Experiments 1 and 2, social behaviors were recorded by camera and scored manually. For the social interaction test, the number and duration of active social interactions were measured, including sniffing, grooming, following, mounting, wrestling, jumping on, and crawling over/under the partner rat (File and Hyde, 1978). For the social choice test and social conditioned place/cue procedure, time spent in chambers was measured from when the nose-point of a rat crossed through an archway. For the social conditioned cue preference test, approach to the star-cue was scored when the rat either touched, sniffed, or gnawed on it.

## Results

### Experiment 1: Sign-trackers but not goal-trackers attribute incentive-motivational value to a social-related cue, and they show conditioned place preference and aversion, respectively

Rats underwent seven daily sessions of PCA training and were classified as STs, and IRs, and GTs based upon their average PCA index scores over Sessions 6 and 7. Only STs and GTs were used for further testing. Figure 1 shows that STs and GTs differed in their lever press number (effect of Phenotype; F_(1,29.44)_ = 41.26, p = 4.69 × 10^−7^), latency (effect of Phenotype; F_(1,30.47)_ = 146.07, p = 3.72 × 10^−13^), and probability (effect of Phenotype; F_(1,28.38)_ = 163.75, p = 2.68 × 10^−13^) as well as magazine entry number (effect of Phenotype; F_(1,31.4)_ = 30.37, p = 4.47 × 10^−6^), latency (effect of Phenotype; F_(1,27.61)_ = 31.73, p = 5.18 × 10^−6^), and probability (effect of Phenotype; F_(1,31.15)_ = 48.08, p = 8.67 × 10^−5^). Rats also differed on their PCA index score (data not shown; effect of Phenotype; F_(1,29.1)_ = 193.22, p = 2.11 × 10^−14^).

**Figure 1.**
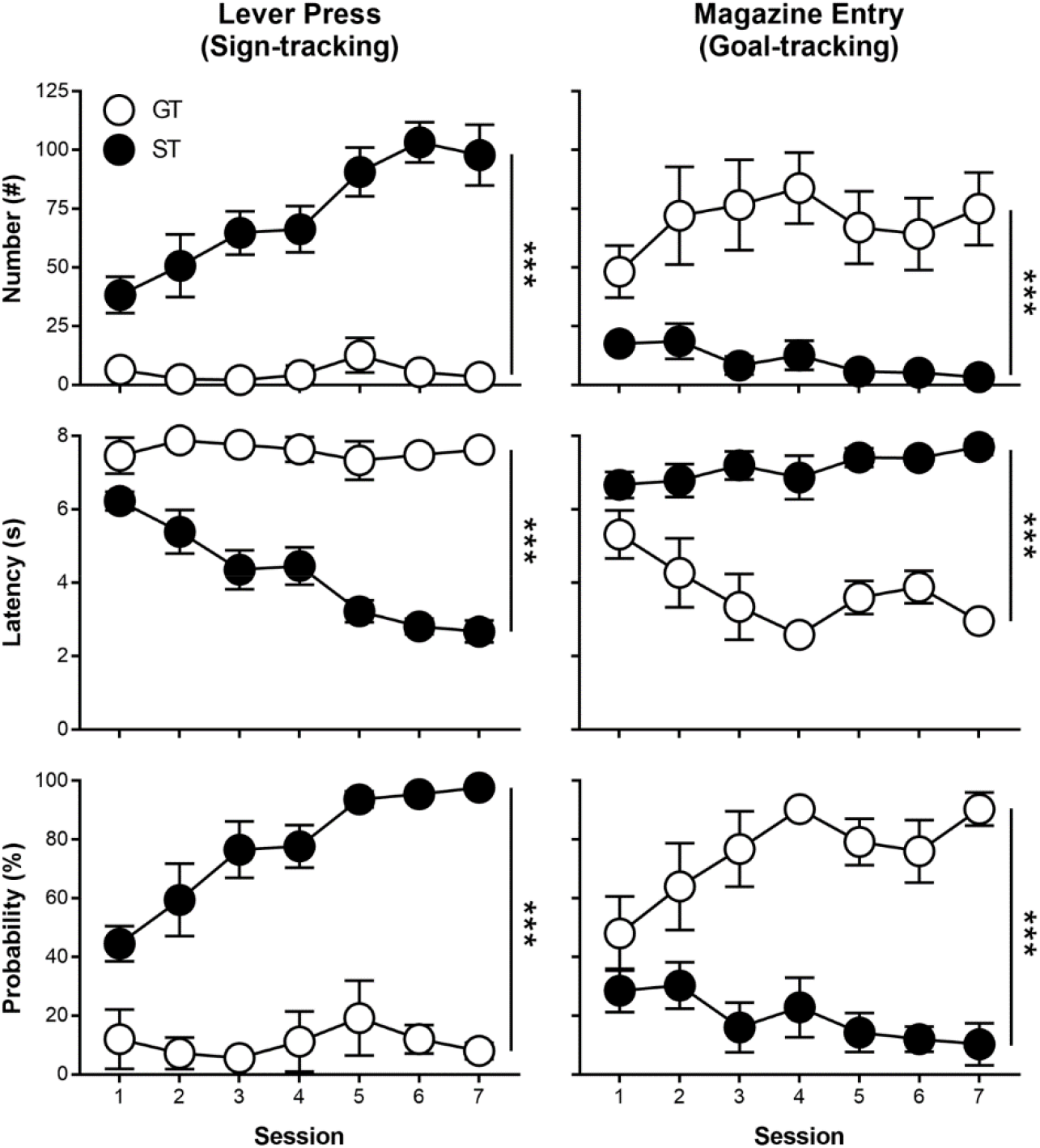
Rats underwent PCA training over seven daily sessions and were classified as sign-trackers (STs), goal-trackers (GTs), or intermediate-responders (IRs) based on their lever press and magazine entry number, latency, and probability during Sessions 6 and 7. Only GTs (n = 5) and STs (n = 7) were used for subsequent testing (social choice and interaction tests) in Experiment 1. Data are presented as mean ± S.E.M. *** - p < 0.001.

During the social conditioned place and cue preference test (Figure 2*A*), each rat displayed a preference for one of the two chambers (i.e., spending 50% or more time in one chamber, averaged over the two daily habituation sessions); however, there was no difference between the time that GTs or STs spent in the preferred or non-preferred chambers (data not shown; interaction of Phenotype x Chamber; F_(31.68)_ = 0.008, p = 0.93). As previously mentioned, rats were conditioned in the nonpreferred chamber. Following eight daily sessions of conditioning (four conditioned and four nonconditioned), rats underwent a conditioned place preference test and cue test. During the place preference test, paired groups conditioned more than unpaired groups, and STs and GTs differed in their place conditioning (Figure 2*B*; interaction of Phenotype x Group; F_(1,19)_ = 10.39, p = 0.004). Post hoc comparisons revealed that paired STs established a place preference and unpaired STs did not (p < 0.05); in addition, paired STs formed a place preference more than paired GTs (p < 0.001). Moreover, paired GTs, but not unpaired GTs, formed a place aversion (p < 0.05). During the cue test, paired groups approached (sign-tracked towards) the star-cue more than unpaired groups, and STs and GTs differed in the number of approaches (Figure 2*C*; interaction of Phenotype x Group; F_(1,18)_ = 5.69, p = 3.01 × 10^−4^). Post hoc comparisons revealed that paired STs sign-tracked to the star-cue more than unpaired STs (p < 0.001) and paired GTs (p < 0.001).

**Figure 2.**
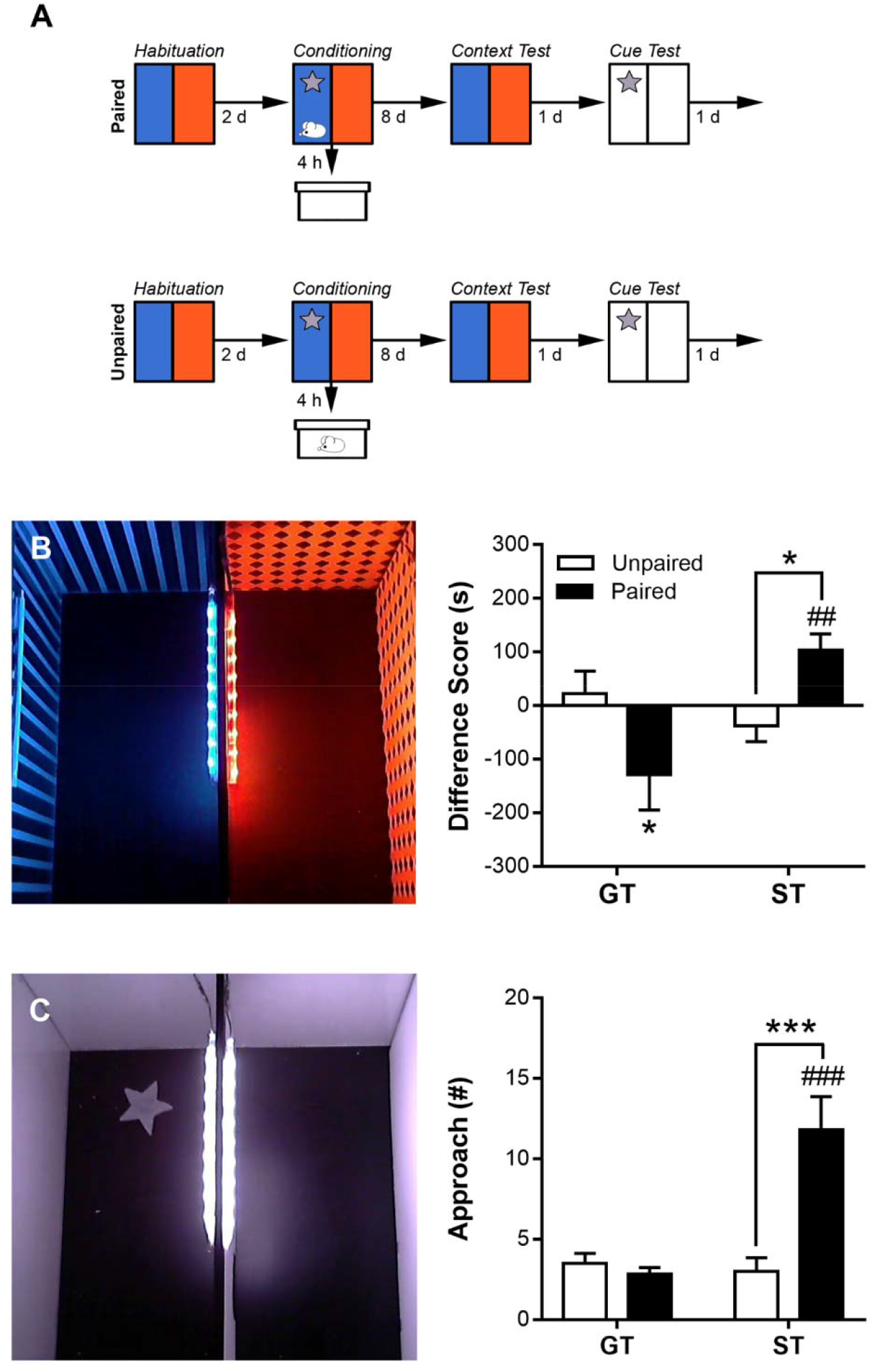
In Experiment 1, seven days after PCA training, sign-trackers (STs) and goal-trackers (GTs) underwent a (A) social conditioned place/cue preference procedure. Following two days of habituation and eight days of conditioning (with the context and a star-cue), paired and unpaired rats underwent a (B) context test and (C) cue test. A difference score (time spent in the social interaction chamber – time spent in the empty chamber) was calculated to measure preference for the context test. Approach to the star-cue was calculated to measure sign-tracking for the cue test. Data are presented as mean + S.E.M. * - p < 0.05, *** - p < 0.001, within-subjects comparison; ## - p < 0.01, ### - p < 0.001, between-subjects comparison.

### Experiment 2: Sign-trackers but not goal-trackers show social novelty-seeking behavior and increased social interaction

Rats underwent seven daily sessions of PCA training and were classified as STs, IRs, and GTs based upon their average PCA index scores over Sessions 6 and 7. Only STs and GTs were used for further testing. STs and GTs differed in their lever press number (data not shown; effect of Phenotype; F_(1,12.95)_ = 59.06, p = 3.54 × 10^−6^), latency (effect of Phenotype; F_(1,15.38)_ = 99.24, p = 4.18 × 10^−8^), and probability (effect of Phenotype; F_(1,15.04)_ = 94.15, p = 7.26 × 10^−8^) as well as magazine entry number (effect of Phenotype; F_(1,13.05)_ = 28.48, p = 1.34 × 10^−4^), latency (effect of Phenotype; F_(1,14.58)_ = 58.01, p = 1.88 × 10^−6^), and probability (effect of Phenotype; F_(1,12.35)_ = 33.32, p = 7.91 × 10^−5^). Rats also differed on their PCA index score (data not shown; effect of Phenotype; F_(1,13.97)_ = 155.84, p = 5.76 × 10^−9^).

Seven days following the last session of PCA training, rats underwent a social choice test, which consisted of three consecutive phases: habituation, sociability, and social novelty. During the habituation phase, neither GTs (Figure 3*A*; effect of Chamber; F_(2,12)_ = 0.43, p = 0.66) nor STs (effect of Chamber; F_(2,18)_ = 1.38, p = 0.28) showed an initial preference for any of the chambers. Next, during the sociability phase, GTs (Figure 3*B*; effect of Chamber; F_(2,12)_ = 39.28, p = 5.42 × 10^−6^) and STs (effect of Chamber; F_(2,18)_ = 109.84, p = 8.19 × 10^−11^) both preferred the chamber containing a partner rat. Post hoc comparisons revealed that both STs and GTs spent more time in the chamber containing a partner than the center (p < 0.001) and empty (p < 0.001) chambers. Finally, during the social novelty phase, STs (Figure 3*C*; effect of Chamber; F_(2,18)_ = 0.14, p = 2.55 × 10^−6^) but not GTs (effect of Chamber; F_(2,12)_ = 0.14, p = 0.87), showed social novelty-seeking behavior. Post hoc comparisons revealed that STs spent more time in the chamber containing a novel partner compared to the center chamber (p < 0.001) and chamber containing the familiar rat (p < 0.001). In addition, post hoc comparisons revealed that GTs spent more time in the chambers containing rats compared to the center chamber (p < 0.01); however, they did not discriminate between the chambers containing the familiar and novel rats (p > 0.05).

**Figure 3.**
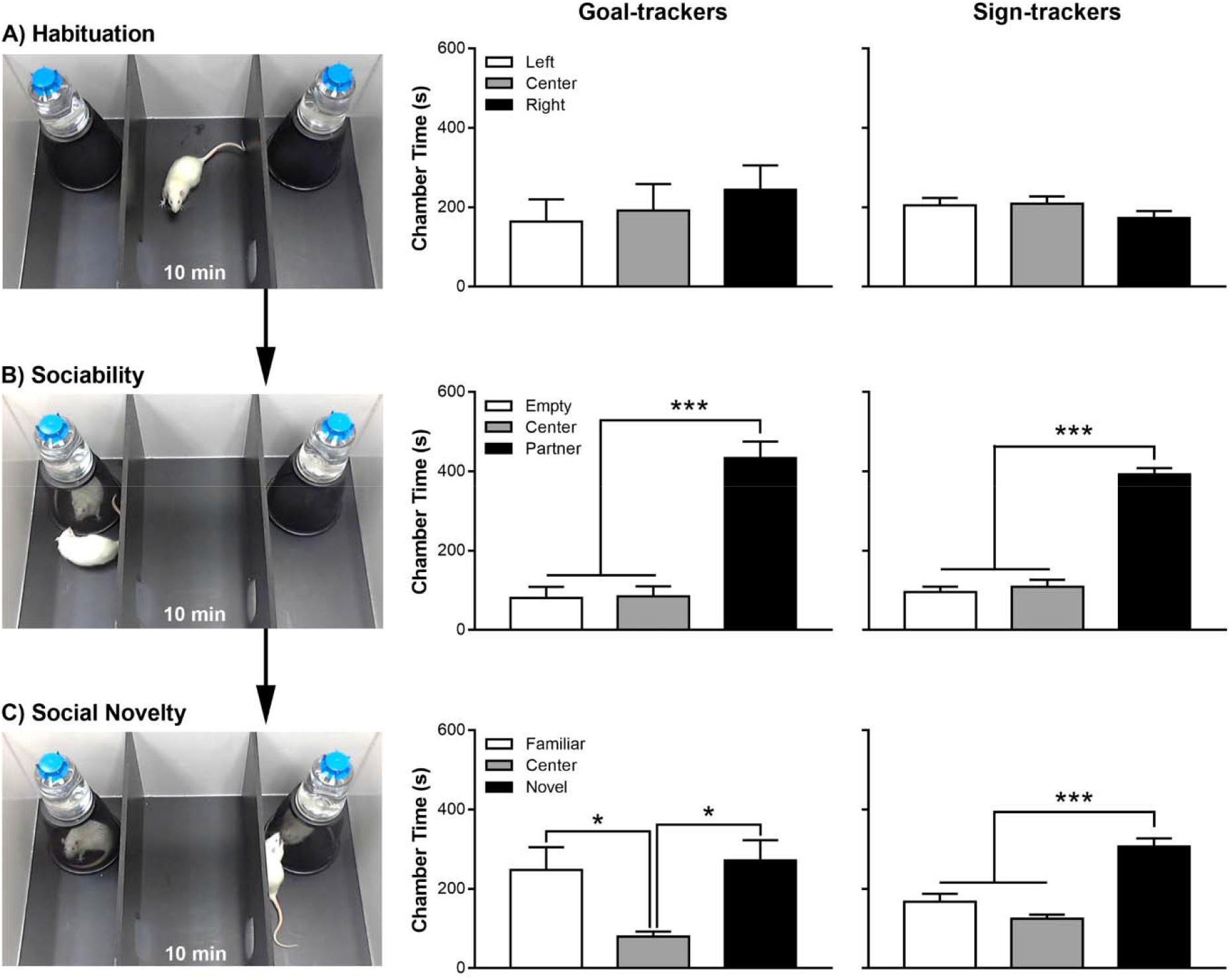
In Experiment 2, sign-trackers and goal-trackers underwent a social choice test that consisted of three consecutive 10-min phases: (A) habituation (exposure to the test arena), (B) sociability (exposure to a partner rat), and (C) social novelty (simultaneous exposure to the now familiar partner rat and a novel partner rat). Data are presented as mean and S.E.M. * - p < 0.05, *** - p < 0.001.

Next, seven days after the social choice test, rats underwent a social interaction test during which the number and durations of active social interactions were measured. STs, compared to GTs, performed more social interactions (Figure 4*A*; effect of Phenotype; t_10_ = - 3.52, p = 0.006) and for longer durations (Figure 4*B*; effect of Phenotype; t_10_ = −3.09, p = 0.011). In addition, STs defecated less than GTs during the social interaction test (Figure 4*C*; effect of Phenotype; t_10_ = 2.59, p = 0.027).

**Figure 4.**
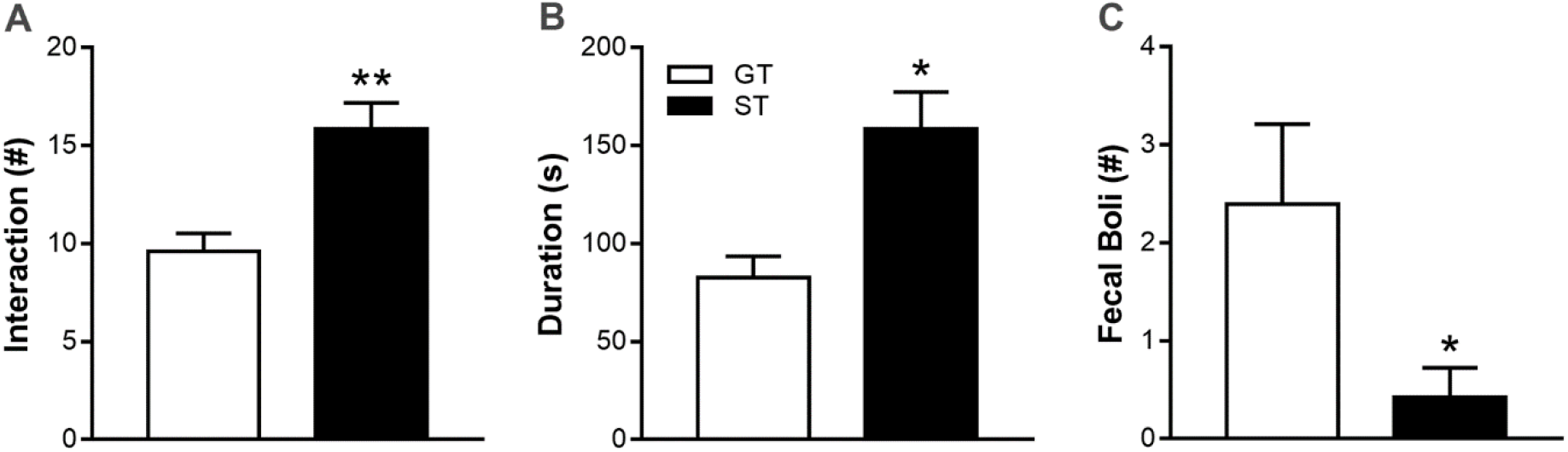
In Experiment 2, sign-trackers (STs) and goal-trackers (GTs) underwent a social interaction test and the (A) number and (B) duration of social interactions as well as (C) fecal boli (a measure of social anxiety-like behavior) were measured. Data are presented as mean and S.E.M. * - p < 0.05, ** - p < 0.01.

### Experiment 3: Baseline plasma OXT levels are not different between GTs, IRs, and STs

Rats underwent seven daily sessions of PCA training and were classified as STs, and IRs, and GTs based upon the PCA index score from Sessions 7. One rat was excluded for not learning any conditioned response (i.e., Session 7 PCA index score = 0). STs, IRs, and GTs differed in their lever press number (data not shown; effect of Phenotype; F_(2,36.16)_ = 41.28, p = 4.64 × 10^−10^), latency (effect of Phenotype; F_(2,37.67)_ = 42.53, p = 2.18 × 10^−10^), and probability (effect of Phenotype; F_(2,37.86)_ = 53.36, p = 9.65 × 10^−12^) as well as magazine entry number (effect of Phenotype; F_(2,39.32)_ = 19.47, p = 1.33 × 10^−6^), latency (effect of Phenotype; F_(2,37.98)_ = 16.85, p = 5.78 × 10^−6^), and probability (effect of Phenotype; F_(2,36.40)_ = 13.13, p = 5.08 × 10^−5^). Rats also differed on their PCA index score (data not shown; effect of Phenotype; F_(2,36.01)_ = 41.81, p = 4.10 × 10^−10^). Seven days following the last session of PCA training, rats were removed from their home cages and baseline plasma OXT samples were collected. GTs, IRs, and STs did not differ in their baseline levels of OXT (Figure 5*A*; effect of Phenotype; F_(2,25)_ = 0.81, p = 0.46). In addition, baseline levels of OXT did not correlate with PCA index scores (r = −0.22, p = 0.25).

**Figure 5.**
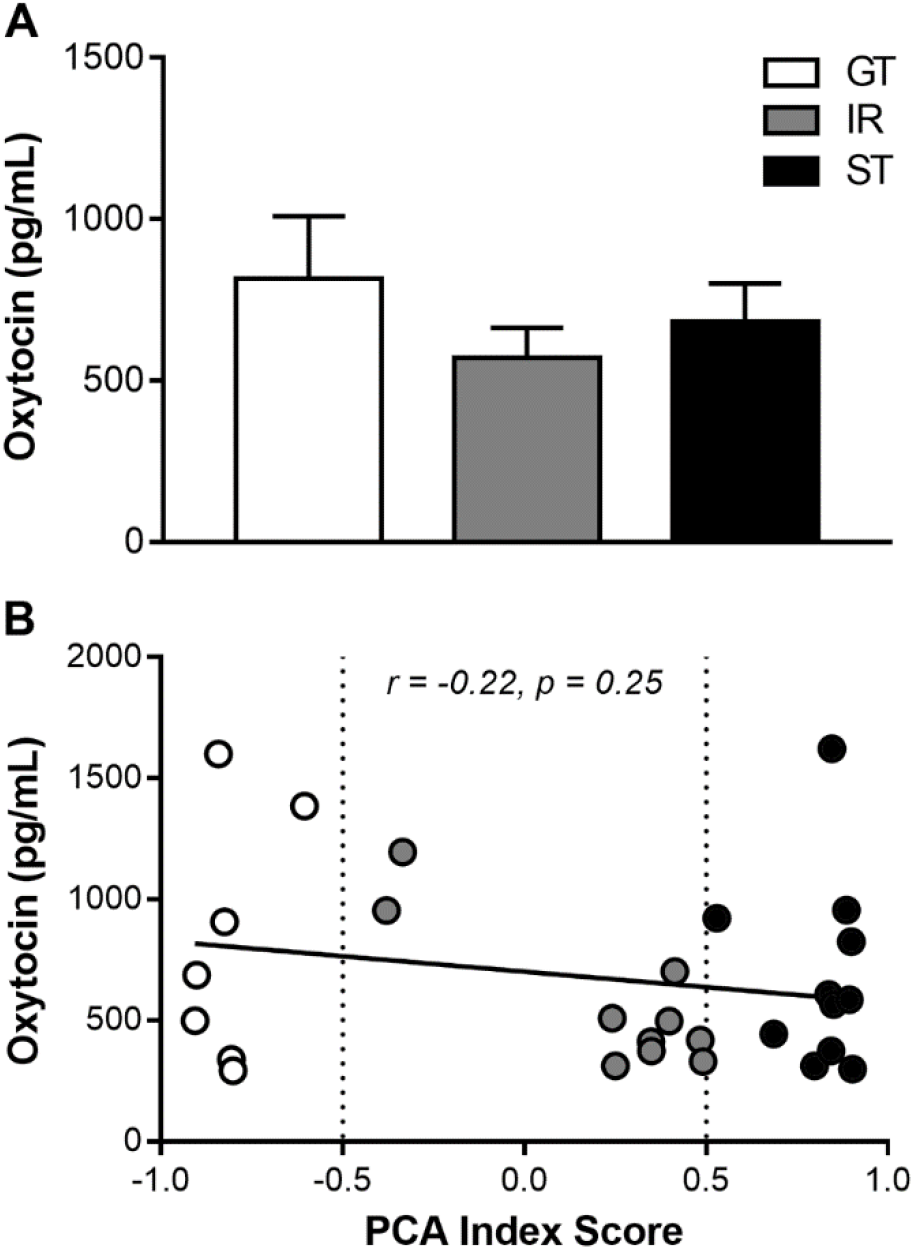
In Experiment 3, baseline plasma oxytocin samples were collected from the home cage seven days after the last Pavlovian conditioned approach (PCA) training session. (A) Plasma oxytocin samples were compared between goal-trackers (GTs), intermediate-responders (IRs), and sign-trackers (STs). In addition, (B) plasma oxytocin levels were correlated with PCA index scores (averaged between Sessions 6 and 7). Data are presented as mean and S.E.M.

## Discussion

In the present study, we demonstrated convergent individual variation in the attribution of incentive-motivational value to a food cue and to a social cue. In other words, rats that sign-track towards a food-related CS also sign-track towards a social cue. In addition, we demonstrated that STs display prosocial behaviors (i.e., social conditioned place preference, social novelty-seeking and increased social interaction) while GTs exhibit antisocial behaviors (e.g., social conditioned place aversion, absent social novelty-seeking, and anxiety-like behavior during social interaction). Finally, PCA phenotypes did not differ in basal plasma levels of OXT, and OXT did not correlate with PCA index scores. Taken together, these results show that (1) social cues can promote sign-tracking behavior, (2) the attribution of incentive-motivational value to reward cues contributes to individual variation in social behaviors, and (3) basal OXT levels are not different between PCA phenotypes.

The present study is the first to show sign-tracking to a nonsexual, social cue. Previously, it has been demonstrated that Japanese quail sign-track towards sexual reward-related cues (Burns and Domjan, 1996). Humans also appear to sign-track toward sexual cues (Kimura et al., 1990). Despite their similarities, however, sexual and nonsexual social rewards have distinct valences and motivational properties (Trezza et al., 2011). Also, our results demonstrate that STs can attribute incentive-motivational value to more than one CS (e.g., a lever-CS predicting food delivery and a star-CS predicting social interaction), and behaviors related to sign-tracking (increased novelty-seeking and CPP) are consistent across different stimuli and procedures (Dickson et al., 2015, Meyer et al., 2012). Finally, this study is the first to demonstrate individual variation in social behaviors in an outbred rodent population. Previously, it has been demonstrated that there are strain differences in conditioned place preference/aversion, social interaction, and social novelty-seeking in rodents (Knoll et al., 2018, Moy et al., 2009, Pinheiro et al., 2016, Ryan et al., 2008); however, all these studies were performed in inbred populations.

Sign-tracking to food and drug-related cues requires DA signaling in the NAc (Flagel et al., 2011, Saunders and Robinson, 2012, Saunders et al., 2013). Because the NAc encodes reward and motivation for social cues in an analogous manner to food- and drug-related cues, it is highly likely that sign-tracking to social cues—like drug- and food-related cues—is DA-dependent in the NAc (Rademacher et al., 2017). In addition, DA signaling in the NAc encodes and is sufficient to regulate social behavior (Gunaydin et al., 2014), suggesting that individual differences in NAc DA signaling may underlie differences in social behavior observed in the present study. For example, increased DA signaling in the NAc of STs during social experiences can explain the formation of CPP (Shippenberg and Herz, 1987), increased social interaction (Manduca et al., 2016), and social novelty-seeking (De Leonibus et al., 2006). Conversely, decreased DA signaling in the NAc of GTs during social experiences can also explain the formation of conditioned place aversion (Shippenberg and Herz, 1987) and decreased social interaction (Manduca et al., 2016).

OXT is a neuropeptide that has a documented role in promoting social behavior and reward by activating dopaminergic pathways in the mesolimbic reward system in response to social experiences and cues (Liu and Wang, 2003, Melis et al., 2007, Rademacher et al., 2017). For example, OXT promotes attention to and processing of social cues (Groppe et al., 2013, Pfundmair et al., 2017), enhances social interaction (Witt et al., 1992), facilitates social discrimination and novelty-seeking (Benelli et al., 1995, De Dreu et al., 2015, Popik et al., 1992), produces CPP (Liberzon et al., 1997), and is necessary for social memory formation (Ferguson et al., 2000). In addition, OXT activates a social learning circuit—the prefrontal cortex, lateral septum, amygdala, ventral hippocampus, NAc, and ventral tegmental area (Dumais et al., 2017, Ferris et al., 2015, Ophir, 2017)—that overlaps with the “motive circuit” underlying sign-tracking behavior (Flagel et al., 2011, Yager et al., 2015). In the present study, baseline plasma levels of OXT did not differ between GTs, IRs, and STs; in addition, plasma OXT levels did not correlate with PCA index scores. It is possible, however, that OXT plasma levels are different immediately following social experiences or presentation of social cues. Future studies should investigate the release of OXT, both centrally and peripherally, in response to social experiences and cues.

In addition to OXT, other neurohormones might contribute to social sign-tracking and individual differences in social behaviors. For example, dynorphin and kappa opioid receptor (KOR) signaling is implicated in the regulation of DA signaling, negative affective states, cue-directed behavior, and social memory formation (Bilkei-Gorzo et al., 2014, Schank et al., 2012, Tejeda and Bonci, 2018). Relevant to the present study, KOR signaling in DA terminals originating from the VTA and terminating in the NAc contribute to conditioned place aversion, social avoidance and decreased social interaction in rodents (Brain et al., 1985;Chefer et al., 2013, Robles et al., 2014, Tejeda et al., 2013). Because KOR agonism decreases DA release in the NAc (Lindholm et al., 2007) and produces negative affective states, it is possible that increased KOR signaling is responsible for the antisocial behaviors in GTs (e.g., conditioned place aversion, decreased social interaction, and lack of social novelty-seeking, and anxiety-like behavior during social interaction). Conversely, decreased KOR signaling surrounding social experiences might permit STs to attribute incentive-motivational value to social cues.

The results from the present study have important implications for the treatment of addiction, because social cues and context are believed to contribute to the pathophysiology of drug addiction (Leshner, 1997). For example, social cues (Breiner et al., 2018, Tsai et al., 2009) and contexts (Garcia-Rodriguez et al., 2011, Huh et al., 2016, Nees et al., 2012) impact reward anticipation and craving. Furthermore, social pressure from peers can induce cravings and promote drug use (Epstein et al., 2009, Lee et al., 2008), and drug users show greater striatal activation during peer conformity to social information (Gilman et al., 2016, Gilman et al., 2016). Conversely, social rewards can also acquire enough motivational value to successfully compete with addictive substances and promote voluntary abstinence from drug self-administration (Venniro et al., 2018). One of the most effective psychotherapies for addiction is coping and social skills training CSST), which teaches social strategies for navigating social interactions, addressing interpersonal problems, and managing craving in response to social/drug-related contexts and cues (Monti and O’Leary, 1999). In addition, new behavioral interventions for addicted patients are centered around social networks (Day et al., 2013), which are an important modulator of drug use and relapse (Mundt, 2011). A better understanding of individual variation in social sign-tracking and social behaviors can aid the development and refinement of therapeutic interventions aimed at restoring a healthy balance between prosocial behaviors and the pursuit of nonsocial rewards in patients with addiction and other related disorders.

## Declaration of conflicting interests

The authors declared no potential conflicts of interest with respect to the research, authorship, and/or publication of this article.

## Acknowledgements

This work was funded by the University of Michigan Department of Psychiatry (U032826 [JDM]) and Rackham Graduate School (Predoctoral Fellowship, CJF), the Department of Defense (DoD) National Defense Science and Engineering Graduate (NDSEG) Fellowship (CJF), and the National Institute on Drug Abuse (NIDA; K08 DA037912-01 [JDM]; T32 DA007281 [CJF]).

## Abbreviations

CS: conditioned stimulus;
DA: dopamine;
ELISA: enzyme-linked immunosorbent assay;
GT: goal-tracker;
KOR: kappa opioid receptor;
NAc: nucleus accumbens;
OXT: oxytocin;
PCA: Pavlovian conditioned approach;
ST: sign-tracker;
US: unconditioned stimulus;
VI: variable interval

